# Resting state MEG oscillations show long-range temporal correlations of phase synchrony that break down during finger-tapping

**DOI:** 10.1101/014159

**Authors:** Maria Botcharova, Luc Berthouze, Matthew J. Brookes, Gareth R. Barnes, Simon F. Farmer

## Abstract

The capacity of the human brain to interpret and respond to multiple temporal scales in its surroundings suggests that its internal interactions must also be able to operate over a broad temporal range. In this paper, we utilise a recently introduced method for characterising the rate of change of the phase difference between MEG signals and use it to study the temporal structure of the phase interactions between MEG recordings from the left and right motor cortices during rest and during a finger-tapping task. We use the Hilbert transform to estimate moment-to-moment fluctuations of the phase difference between signals. After confirming the presence of scale-invariance we estimate the Hurst exponent using detrended fluctuation analysis (DFA). An exponent of >0.5 is indicative of long-range temporal correlations (LRTCs) in the signal. We find that LRTCs are present in the *α*/*μ* and *β* frequency bands of resting state MEG data. We demonstrate that finger movement disrupts LRTCs correlations, producing a phase relationship with a structure similar to that of Gaussian white noise. The results are validated by applying the same analysis to data with Gaussian white noise phase difference, recordings from an empty scanner and phase-shuffled time series. We interpret the findings through comparison of the results with those we obtained from an earlier study during which we adopted this method to characterise phase relationships within a Kuramoto model of oscillators in its sub-critical, critical and super-critical synchronisation states. We find that the resting state MEG from left and right motor cortices shows moment-to-moment fluctuations of phase difference with a similar temporal structure to that of a system of Kuramoto oscillators just prior to its critical level of coupling, and that finger tapping moves the system away from this pre-critical state towards a more random state.

## 1 Introduction

The human MEG is a rich time-varying signal that provides an important window into ongoing brain processing [1–4]. In the frequency domain the MEG is characterised by spectral peaks overlaying 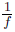 noise, such that its power at each frequency diminishes in an inverse relationship [3]. The power of the spectral peaks varies with sensor location and task performed, but in general terms there are peaks of spectral power in the *θ*, *α*/*μ*, *β* and *γ* bands [5]. Underlying these peaks is oscillatory neuronal activity due to network synchronisation [3]. In the motor system there is particular interest in the behaviour of spectral peaks in the *α*, *μ* and *β* bands whose oscillatory activity dominates in resting state MEG recorded over the sensory-motor cortex and which is diminished by transient movement of the contra-lateral muscle groups [6, 7]. During bimanual movement and motor learning there are changes in EEG synchronisation between different motor areas including left and right motor cortices [8]. During sustained muscle contraction there is *β*-band synchronisation between motor cortex EEG/MEG and the contralateral EMG [9–11]. Importantly neural synchronization is weak and varies with time even during a sustained task. These fluctuations of synchrony can be detected using a variety of analytical techniques: time-varying coherence, wavelet coherence and optimal spectral tracking [12, 13]. The observation that oscillatory brain signals may come in and out of synchronisation with each other spontaneously and with task, and idea that synchronisation lends salience to a signal, has led to the hypothesis of ‘communication through coherence’ (CTC). Implicit is the idea that the variability of the effective coupling is in itself of significance [12].

Because neural signals exhibit moment-to-moment fluctuations in their amplitude and phase it is possible to characterise the temporal correlations within such time series. The order within amplitude fluctuations of bandpass-filtered MEG and EEG time series has been previously quantified through estimation of the Hurst exponent [14, 15]. The exponent of a power law relationship between scale and fluctuation magnitude within that scale characterises the presence of long-range temporal correlations in the time series. An exponent of 0.5 indicates white noise with no temporal correlation, whereas exponents of >0.5 or <0.5 indicate either long-range or an anti-correlated time series, respectively. These data have been interpreted within the framework of the ‘critical brain hypothesis’ in which the presence of spatio-temporal correlations in brain signals at single unit level, at LFP level and at surface signal level (such as MEG and EEG oscillations) are thought to reflect a system operating close to a critical regime [16–18]. Given the importance of neural synchrony and its fluctuation for communication between brain regions, it is of interest to ask whether in human brain signals there is evidence that interareal synchronization exists within or close to a critical regime [19].

In this paper we explore whether there are LRTCs in the moment-to-moment fluctuations of the MEG phase difference between different brain regions and whether, if present, these are disrupted by finger movement. Using resting state human MEG data, we determine first if there is power law scaling and if so we derive the DFA exponent of the fluctuations in the rate of change of phase difference between two simultaneously recorded MEG signals obtained from left and right primary motor cortices. We next explore the effects of bilateral muscle activation (finger tapping) on the MEG phase difference scaling and the associated DFA exponent. Finger movement is known to disrupt sensori-motor cortex oscillations [6] and may also act as feedback perturbation of the type that has been shown to reduce LRTCs in the amplitude envelope of the EEG and MEG [15]. Here we use finger tapping for its potential to disrupt the temporal organisation of left and right MEG phase differences in the resting state. We show that in the resting state the moment-to-moment fluctuations of phase difference between the left and right motor cortex MEG time series has both power law scaling and an estimated DFA exponent >0.5 in the *α*/*μ* and *β* frequency bands, indicative of long-range temporal correlations within moment-to-moment fluctuations of phase difference. Finger tapping movements, when compared to the resting state, are associated with a significantly lower DFA exponent in the MEG moment-to-moment fluctuations of phase difference in the *α*/*μ* and *β* frequency bands. We validate our results by applying the same analysis to data with Gaussian white noise phase difference, recordings from an empty MEG scanner and phase-shuffled time series. The demonstration of the presence of scaling with exponent values consistent with the presence of long-range temporal correlations in this data provides an important connection between the critical brain and neural synchrony paradigms.

## 2 Material and methods

A key novel component of this work is our application of a method that allows detection of the presence of long-range temporal correlations (LRTCs) in the moment-to-moment fluctuations of phase difference between two neurophysiological time series. In Sections 2.1–2.3, we first detail the steps involved in the extraction of the rate of change of the phase difference time series from the neurophysiological data and then briefly summarise the two techniques used to rigorously assess the presence of LRTCs, along with statistical testing.

In Section 2.4 we give details of the subjects and data recordings used to collect the data. Finally, Section 2.5 describes the control time series used to validate the results, including analysis of synthetic noise and empty scanner and phase-shuffled MEG data.

The overall methodology for the extraction and analysis of the neurophysiological signals used in this manuscript follows that proposed by the authors in [20] with modifications suitable to its application to MEG data. The methodology is illustrated in a step-by-step fashion in Figure 1. Below, we detail each step in turn.

**Figure 1:**
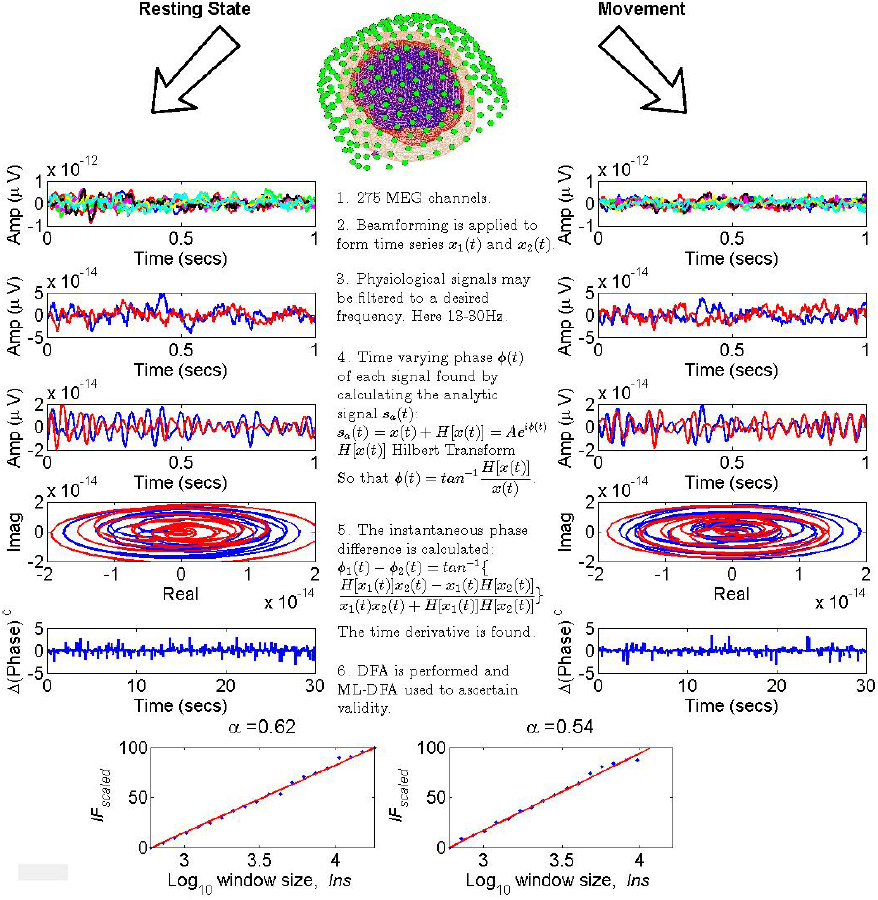
Step-by-step illustration of the method. Each numbered step corresponds to a pair of plots in the diagram for resting state and movement data. The time series in the panels on the left are taken from resting state MEG data, and those in the panels on the right are from human MEG data recorded while the subject was tapping both index fingers simultaneously. The red and blue colours in steps 2-4 correspond respectively to the MEG time series for the left and the right motor cortices. The time segment of each time series remains consistent from one step to the next. Step 1) Data from the full set of 275 MEG channels is considered. Step 2) The data is beamformed to obtain two time series corresponding to the left and right motor cortices respectively. Step 3) Each time series is bandpass-filtered to a frequency of choice as described later in the text. Step 4) The analytic signal *s*_*a*_(*t*) for each time series is found and plotted on a phase space diagram on axes representing the real and imaginary parts of *s*_*a*_(*t*). Step 5) The phase difference between the two time series is found by taking its arctangent. The first time derivative of this phase difference is then taken to produce the time series plotted. Step 6) DFA is applied to obtain an exponent and ML-DFA is used to ascertain whether ^5^this exponent is valid. For the data shown here, both DFA plots were judged to be linear. The brain image was generated using SPM8 [21].

### 2.1 Extraction of the phase difference time series

#### 2.1.1 Beamforming

The MEG data were initially processed using a scalar linearly constrained minimum variance beamformer [22] in order to estimate timecourses of source strength at locations of interest in the left and right motor cortices. The forward models, covariance matrices and calculation of beamformer weights are described in [23]. The weights for left and right motor cortex were based on those contralateral voxels showing the largest change in *β*-band power for right and left hand movement respectively [7]. This was done individually for each of the 7 subjects.

#### 2.1.2 Bandpass Filtering

The use of the Hilbert transform (see Section 2.1.3) requires a narrowband signal for the phase and amplitude components of the signal to be straightforward to separate [24, 25]. We filtered the data with a finite impulse response (FIR) filter. The filter order of the FIR filter was set to include three cycles of the lower-frequency component of the frequency band. A similar approach has been used in many recent studies, e.g., [14].

The frequency bands were selected as 2Hz intervals spanning the full range of frequencies to 4.5-39.5 Hz following a technique used in Klimesch et al. [26]. In this paper, the authors identify peaks in the power spectrum, which are then used to define frequency bands …, f-2 to f, f to f+2, f+2, f+4, … Here, the resting state power spectra have two peaks in two frequency bands one at 10.5 Hz in the *α*/*μ* frequency range and another at 21.5 Hz in the *β* frequency range (see Section 3.1). For this reason the time series were filtered in bands 4.5-6.5 Hz, 6.5-8.5 Hz, …, 14.5-16.5 Hz in the *θ*, *α*/*μ* frequency range, and in bands 15.5-17.5 Hz, …, 37.5-39.5 Hz in the *β* and *γ* frequency ranges.

#### 2.1.3 Phase difference

The phase relationship *ϕ*_1_(*t*) – *ϕ*_2_(*t*) between two different time series *s*_1_(*t*) and *s*_2_(*t*) was calculated using the respective Hilbert transform of the signals *H*[*s*_1_(*t*)] and *H*[*s*_2_(*t*)] [27]:

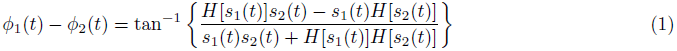

The time series *ϕ*_1_(*t*) – *ϕ*_2_(*t*) is an unbounded process because *ϕ*_1_(*t*) and *ϕ*_2_(*t*) themselves are unbounded as long as the signals *s*_1_(*t*) and *s*_2_(*t*) continue to evolve as time increases. As we use detrended fluctuation analysis (DFA), see Section 2.2.1, to assess the presence of long-range temporal correlations and DFA in its standard form assumes a bounded signal, in this paper, we characterise phase synchronisation in terms of the rate of change (first time derivative) of the phase difference time series *ϕ*_1_(*t*) – *ϕ*_2_(*t*).

### 2.2 Characterisation of the LRTCs

Long-range temporal correlations (LRTCs) are present in a time series when its autocorrelation function, *R*_*ss*_(*τ*), has a decay that takes the form of a power law, such that *R*_*ss*_(*τ*) ∼ *Cτ*^−*α*^ where *τ* is the time lag between observations and *α* is the exponent of the power law. One approach to determining this exponent is to estimate the Hurst exponent, which directly relates to *α* through *α* = 2 - 2H [28, 29], using Detrended Fluctuation Analysis [30].

#### 2.2.1 Detrended Fluctuation Analysis (DFA)

To calculate the DFA exponent, the time series (in this case the rate of change of phase difference between the two MEGs) is first detrended and then cumulatively summed. The root mean square error is then calculated when this signal is fitted by a line over different window sizes (or scales). Extensions of the technique can be used to fit any polynomial to each window, however, here we only consider linear detrending. If the time series is self-similar, there will be power law scaling between the residuals (or detrended fluctuations) and the window sizes. In log space, this power law scaling yields a linear relationship between residuals and window sizes, the so-called DFA fluctuation plot, and the DFA exponent H is obtained using least squares linear regression. A DFA exponent in the range 0.5<H<1 indicates the presence of long-range temporal correlations. An exponent of 0<H<0.5 is obtained when the time series is anti-correlated, *H*=1 represents pink noise. Gaussian white noise has an exponent of *H*=0.5.

#### 2.2.2 Data Stitching

In our application of DFA to MEG data, we used 20 window sizes with a logarithmic scaling and a minimum window of 1 second, guaranteeing several cycles of the slowest oscillation. To improve the estimate of the DFA, trials during which a given subject was performing the same task were stitched together to produce a single, longer time series for each subject (see below for more detail). We used a maximum window size of 30 seconds, which is the length of a single trial, and equivalent to N/10 where N is the length of the stitched time series.

Stitching is a commonly used step and does not affect the numerical results obtained by DFA if the window sizes are set appropriately [31]. Since the windows are chosen on a logarithmic scale, there will be some windows at some window sizes in which distinct trials of data overlap. This is acceptable for longer data sets (as formed by stitching here) because at least half of the windows will always be contained within a continuously recorded segment of data.

When stitching is applied to time series, it is done between steps 5 and 6 of our method (Figure 1). Namely, the dividing and stitching together of trials should be performed after beamforming, filtering and calculating the phase difference. This minimises the chance of edge artefacts from the Hilbert transform [32, 33].

#### 2.2.3 Assessing the validity of DFA

An interpretable DFA exponent is obtained when the log-log plot of fluctuation size against window size is truly linear, i.e., there is power law scaling (see Section 2.2.1). Since there is no *a priori* means of confirming that a signal is indeed self-similar, an exponent can always be obtained even though the DFA fluctuation plot may not necessarily be linear – the only certainty being that it will be increasing (albeit not necessarily monotonically so) with window size.

We used the technique described in [34] to determine whether each DFA fluctuation plot is well-approximated by a linear model. This is a heuristic technique, which has been tested extensively on known time series in order to verify the robustness of its conclusions [34]. The technique first fits the DFA fluctuation plot with a number of candidate models. These models (polynomial, root, logarithmic, exponential and spline with 2 to 4 sections) were chosen based on published characterisations of DFA fluctuation plots obtained with timeseries of known properties, e.g., with superposed trends or with noise. The number of parameters in these models range from 2 for the linear model to 8 for the four-segment spline model. The technique then maximises a function that is similar in form to a log-likelihood:

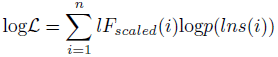

where *lF*_*scaled*_ are normalised fluctuation magnitudes, i indexes the windows, and

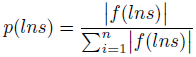

where *f*(*lns*) is the fitted model. The fits of all models are compared using the Akaike Information Criterion (AIC), which discounts for the number of parameters needed to fit the model. The DFA exponent is accepted as being valid only if the best fitting model (that with the lowest value of AIC) is the linear model. Only those time series that were not rejected for not being linear contributed to our results.

### 2.3 Statistical tests of MEG data analysis

We corrected for multiple comparisons using a permutation test in each of the three bands (*α*/*μ*, *β* and *γ*) [35]. In particular, we found the mean of the difference in DFA exponent between the resting state and finger tapping task for each subject and each sub-band of the *α*/*μ*, *β* and *γ* bands. We then swapped the DFA exponents for resting state and finger tapping task in some of the subjects. This could be done in 2^7^ = 128 ways, and we performed each one. For each permutation we found the mean of the difference in DFA exponent between the tasks defined by the shuffled labels. We then took the maximum of these mean differences across all sub-bands. This gave a null distribution of the differences in DFA exponent between tasks that we would expect by chance. We repeated this procedure to create separate significance thresholds for each of three bands of interest (*α*/*μ*, *β* and *γ*).

### 2.4 Participants

Seven healthy right handed subjects took part in the MEG experiments. The study was approved by the University of Nottingham Medical School ethics committee. Details of the experimental procedure are described in Brookes et al. [23]. In brief, MEG data were recorded using the third order gradiometer configuration of a 275 channel CTF MEG system at a sampling rate of 600 Hz. The scanner was housed inside a magnetically shielded room (MSR) and a 150 Hz low pass anti-aliasing hardware filter was applied. All subjects underwent a single experiment which comprised 10 trials of rest and movement. A single trial comprised 30 s during which both left and right index fingers were moved together, and 30 s of rest. The movement itself comprised abductions and extensions of the index fingers. The motor task was cued visually via projection through a waveguide in the MSR onto a back projection screen located 40 cm in front of the subject.

During data acquisition the location of the subject’s head within the scanner was measured by energizing coils placed at 3 fiducial points on the head (nasion, left preauricular and right preauricular). If any subject moved more than 5 mm during the experiment, data from that subject was discarded. Following data acquisition, the positions of the coils were measured relative to the subject’s head shape using a 3D digitizer (Polhemus isotrack). An MPRAGE structural MR image was acquired using a Philips Achieva 3T MRI system (1 mm^3^ isotropic resolution, 256×256×160 matrix size, TR=8.3 ms, TR=3.9 ms, TI=960 ms, shot interval=3 s, FA=8° and SENSE factor=3). The locations of the fiducial markers and MEG sensors with respect to the brain anatomy were determined by matching the digitized head surface to the head surface extracted from the 3T anatomical MRI.

### 2.5 Control Data

In order to search for artefactual effects, we applied our method to three different classes of both synthetic and experimental control time series.

#### 2.5.1 Synthetic data with noise phase difference

Two time series *s*_1_(*t*) and *s*_2_(*t*) were constructed such that their phase difference was white Gaussian noise with a known DFA exponent of 0.5 [36]. The signals were calculated by 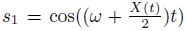 and *s*_2_ = 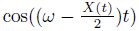. Here, *ω* = 1 and was included because white noise time series alter sharply at each new innovation and the Hilbert transform can produce artefacts when applied to such data. This situation does not arise in physiological data because the sampling frequency with which physiological data are recorded yields a smooth signal.

Seven pairs of time series were created to match the 7 human subjects. Each time series consisted of 10 segments of *X*(*t*) of length 30 seconds which were stitched together to construct the two signals *s*_1_(*t*) and *s*_2_(*t*) (see Section 2.2.2 for more detail). These signals were not beamformed because there was no possibility of unwanted noise or crosstalk. Window sizes used for application of DFA were identical to those used for physiological data, namely logarithmically spaced windows with minimum 1 second and maximum 30 seconds. Although there is no sampling frequency associated with noise simulations, the minimum window was set to 600 time samples for consistency.

The difference between the phases of the two signals *s*_1_ and *s*_2_ is precisely *tX*(*t*) yielding a first time derivative of *X*(*t*). On applying DFA, we would therefore expect the prescribed exponent to be recovered if no bandpass filter was applied. This control data therefore makes it possible to quantify and rule out any artefact potentially introduced by filtering.

#### 2.5.2 Scanner Noise

We analysed 7 data sets recorded from an empty MEG scanner. Time series were obtained from sensors corresponding to the approximate locations of the right and left motor cortices, with a sampling frequency of 600Hz. These continuous time series of length 300 seconds were divided into 10 segments of length 30 seconds each, and reordered randomly to replicate the stitching procedure applied to MEG data. This reordering was performed after step 5 of our method (see Figure 1 and Section 2.2.2 for more detail), which ensured consistency with analysis of the human MEG data. DFA exponents were then obtained for the 7 time series pairs, and checked for validity using ML-DFA. Window sizes used for DFA were identical to those used for the physiological data.

#### 2.5.3 Statistical Tests

The DFA exponents of the synthetic pairs of time series constructed in Sections 2.5.1 and 2.5.2 were tested for normality using a one-sample Kolmogorov-Smirnov test, for each frequency band individually as for MEG data. These tests showed that a null hypothesis of normally distributed DFA exponents could not be rejected for any synthetic time series pair or frequency band where more than one DFA exponent was judged to be valid by ML-DFA (p-values not shown). This was confirmed using the Jaque-Bera test for normality (data not shown). As normal distributions could not be rejected, DFA exponents obtained from the analysis of the above time series were tested for statistical difference with exponents obtained for resting state human MEG data using a Student’s t-test (unpaired).

#### 2.5.4 Phase Shuffling

A permutation of the time derivative of the phase difference time series (the time series in step 5 of our methodology in Figure 1) was performed 100 times yielding 100 DFA exponents. These 100 exponents were tested for significant differences with the 7 human MEG data using the technique presented in [37]. The p-value was approximated using

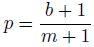

where *b* is the number of DFA exponents obtained from the permuted time series that are greater in size than that of the correctly ordered human MEG data, and *m* is the number of permutations used *m* = 100. This is an approximation to the p-value when the number of possible permutations of the time series is very large, as in this case where the number of possible permutations is 180, 000! (factorial).

## 3 Results

### 3.1 Spectral analysis

We analysed data from the MEG oscillations of the left and right motor cortices during rest and during finger tapping tasks. The power spectra of time series for all 7 subjects were calculated and using pooled power spectral analysis [38] the most prominent spectral peaks were determined (10.5 Hz and 21.5 Hz). As explained in the Methods, these peaks were used to set the boundaries of the ranges of frequency bands used for bandpass-filtering.

### 3.2 Average DFA exponents of rate of change of phase difference compared to shuffled data

Figure 2 shows for 7 subjects the mean±SD valid DFA exponents calculated from the rate of change of phase difference between left and right hemisphere MEGs. The time series were bandpass-filtered in regular 2Hz bands prior to analysis (see Section 2.1.2). As recommended by [7], we state the boundary values of each frequency band. These data are shown for each 2Hz frequency band across the *θ*/*α*/*μ*, *β* and *γ* frequency ranges (Figure 2). Only experiments associated with fluctuation plots judged to show linear scaling using ML-DFA are included in the data (see Appendix Table 2). The results reveal an estimated Hurst exponent for the rate of change of phase difference between left and right motor cortex MEGs that is >0.5 (i.e., with long-range temporal correlations) at rest and during finger tapping in the *θ*/*α*/*μ* and *β* frequency ranges. The mean±SD DFA values for phase-shuffled MEG data are also shown for each frequency band (Figure 2). Phase shuffling preserves the spectral power of the signal but disrupts its temporal relationships. The results of shuffled data show DFA exponents close to 0.5 (i.e., uncorrelated noise) in all frequency bands as expected. Importantly, the resting state DFA values are significantly greater (as assessed by permutation p-values, see Section 2.5.4) than the corresponding shuffled values for in the *α*/*μ* and *β* frequency ranges. Significant differences between finger tapping DFA values and the corresponding shuffled values are only observed in the *α*/*μ* frequency range.

**Figure 2:**
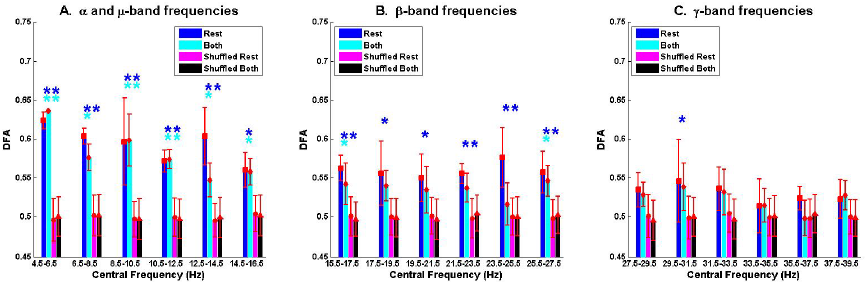
Mean±SD DFA exponents for ’rest’ and ’both’ conditions against the distributions of DFA exponents for phase-shuffled MEG time series for the *θ*/*α*/*μ*, *β* and *γ* frequency ranges (Panels A-C). Mean DFA exponents are in dark blue for resting state and cyan for the finger tapping task, with vertical bars showing standard deviation. The mean DFA exponents of the phase-shuffled resting state are in magenta, with those of the phase-shuffled finger tapping condition in black. Statistical significance for each band and each condition was assessed using permutation p-values calculated with the methodology described in Section 2.5.4 (* for p<.05; ** for p<.01).

### 3.3 Comparison of DFA exponents of resting MEG with finger tapping MEG

The mean±SD differences between the DFA exponents of rest and finger tapping conditions were calculated for each frequency band (Figure 3, blue bars). The corresponding quantities for the phase-shuffled data were also determined (Figure 3, pink bars). Our results show that the average DFA exponents for the resting state are higher than, or comparable to, those obtained from data recorded during the finger tapping task. When correcting for multiple comparisons within each canonical band (see Section 2.3), the difference between these average DFA exponents is shown to be significant in the 12.5-14.5 Hz (p<.05) and 23.5-25.5 Hz (p<.01) frequency bands (Figure 3). Only exponents associated with those fluctuation plots judged to be valid by ML-DFA are included in the data. The number of subjects and shuffled time series contributing to the calculation of each average DFA exponent is listed in Appendix Table 2. In the two frequency bands where statistically significant differences in DFA between rest and movement were seen, there were 6 (out of 7) and 7 (out of 7) valid subject recordings, respectively. We note that these frequency ranges are close, but not identical, to peak power of the MEG in the *α*/*μ* and *β* frequency ranges respectively.

**Figure 3:**
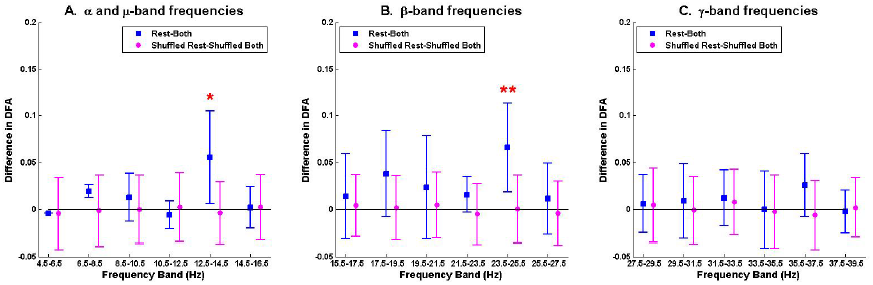
The mean difference of DFA exponents when contrasting rest and movement in the *θ*/*α*/*μ*, *β* and *γ* frequency ranges (Panels A-C). Summary data for the difference between resting state and finger tapping task are shown in blue (rest-both). Data for the corresponding phase-shuffled data are shown in magenta circles (shuffled rest-shuffled both). Statistical significance (corrected for multiple comparisons) is shown by asterisks (* for p<.05; ** for p<.01).

### 3.4 Validation of results

The validity of the >0.5 estimated Hurst exponents detected for the resting state MEG was further assessed for each frequency band through comparing them with (a) synthetic data constructed to yield white Gaussian noise phase difference and (b) empty MEG scanner noise. The number of DFA exponents obtained from analysing phase differences in these control signals was comparable to those of human MEG data in each frequency band (Appendix Table 2). There were significant differences for the resting state data against both MEG scanner noise and Gaussian white noise in the *β*-band (23.5-25.5 Hz). Statistical significance in the *α*/*μ* band was only found against white Gaussian noise in the 10.5-12.5 Hz band and scanner noise in the 12.5-14.5 Hz band. There were no differences in the *γ* frequency range.

## 4 Discussion

### 4.1 Summary

In this paper, it has been shown that resting state human MEG shows power law scaling in the temporal order of the rate of change of the phase difference between left and right motor cortices. The exponent of this scaling was found to be greater than 0.5, which indicates long-range temporal correlations in this measure of synchronisation. Furthermore, it was demonstrated that these long-range temporal correlations are disrupted by finger movement in the 12.5-14.5 Hz and 23.5-25.5 Hz frequency bands.

### 4.2 Methodological considerations

We have applied a novel method to MEG data analysis [20]. It combines a technique for determining the rate of change of the phase difference between two time series with detrended fluctuation analysis (DFA). DFA produces an exponent for a power law relationship between (temporal) scale and fluctuation magnitude within that scale. A further aspect of our methodology is the application of ML-DFA [34] to robustly determine whether there is linear scaling such that a power law in the fluctuation plot exists.

Prior to analysis, the MEG data was beamformed, time series from trials corresponding to the same task for the same subject were stitched together, and the data was then bandpass-filtered. As each of these steps introduces complexities and possible artefacts in the analysis, we discuss these in turn below.

#### 4.2.1 MEG recording

Cross-talk and noise is a potential problem for MEG recordings [39]. The effects of these shortcomings are minimised by identifying the motor cortices individually for each subject using a beamforming technique for β frequencies known to be sensitive to motor tasks [23, 40, 41].

#### 4.2.2 Filtering

The data was bandpass-filtered in bands of 2Hz using a technique from [26]. This method ensures that the frequency bands are sufficiently small to enable the separation of amplitude and phase components of the signal [24] yet sufficiently wide to avoid artefactual coupling [42].

#### 4.2.3 Data stitching

It was necessary to stitch together data for the robust application of the DFA method. A DFA exponent calculated from a time series stitched from shorter segments with comparable exponents of their own will retain validity and will provide a robust estimate of the underlying Hurst exponent if this exponent is in the LRTC range (>0.5) [31]. Further, it has been demonstrated that removing 50% of a LRTC signal and stitching together the remaining parts (even with different standard deviations) does not affect the scaling behaviour of the signal [31].

We note that, since each trial is 30 seconds in length, any periodic artefactual effects of stitching these trials would occur at a frequency of 1/30 Hz. This frequency is below any frequency band used in our analysis. Furthermore, because DFA operates on detrended data and is insensitive to signal amplitude, differences in the mean between segments or the presence of linear trend are unlikely to affect our method [43].

#### 4.2.4 Frequency band dependence

We observed that the number of DFA exponents found to be valid by ML-DFA increased with higher frequency bands whereas their value fell as the frequency band increased, a possible effect of filtering (see Figure 2). To eliminate this effect from the interpretation of our results, we presented the DFA exponents results as differences between rest and finger tapping conditions within the same frequency band such that any artefactual inflation of the DFA exponent as a result of the choice of frequency band in which it was measured, was controlled. Importantly, there was no difference, for any frequency band, in the DFA exponents of the rate of change of phase difference between the two phase-shuffled time series (rest and movement). These time series have the same power spectra as the original (not phase-shuffled) data. We also compared our results to those obtained when the analysis was performed on Gaussian white noise and MEG scanner noise filtered identically to the rest and movement MEG data (see Appendix Table 1). When correcting for multiple comparisons statistically significant increases in DFA exponent values were detected for resting state data in both *α*/*μ* and *β* frequency ranges.

### 4.3 General discussion

In a previous paper [20] we applied this method to phase differences generated by the Kuramoto model with added Gaussian noise. No bandpass filtering was needed as the Kuramoto model produces phase signals. The Kuramoto model has a coupling parameter that can be adjusted. The value of the coupling value at which the Kuramoto system shows a critical transition is known. The critical transition is characterised by a global order parameter which reflects the overall organisation of the system. Our methodology makes it possible to make observations at a pair-wise level, i.e., between individual pairs of Kuramoto oscillator. As individual Kuramoto oscillator pairs become fully synchronised, their phase difference no longer contains moment-to-moment fluctuations and thus power law scaling in the DFA measure is lost [20]. This occurs at and above the critical coupling parameter. When coupling is just below this level, power law scaling exists in the DFA of the rate of change of phase difference time series in the pairs approaching full synchrony. The DFA exponent of oscillator pairs that are not yet fully synchronised is above 0.5, indicating LRTCs for fluctuations of their rate of change of phase difference. We note that in the present study similar DFA values (>0.5) were obtained for the resting state MEG recorded between left and right motor cortices and that the maximal DFA values were frequency band specific. In the Kuramato model with noise, as coupling is decreased further, the DFA exponent of the rate of change of pairwise phase differences decreases towards 0.5. The results obtained from analysis of left and right motor cortex MEG during movement showed retained power law scaling but with DFA exponents close to 0.5 and significantly less than those DFA exponent values obtained in the *α*/*μ* and *β* frequency ranges in the resting state. These results suggest that in the resting state, MEG signals behave as weakly coupled oscillators whose phase difference fluctuations scale and have non-trivial temporal order (estimated Hurst exponent >0.5). This behavior is disrupted by finger movement.

The Kuramoto model makes it possible to take a system of oscillators through the transition from an uncoupled to a synchronised state. In the human brain we are only privy to snapshots of the working system of MEG oscillations. However, by focusing on the fluctuations of the rate of change of phase difference between brain areas we can begin to understand the interaction of brain oscillations through comparison of the results of this analysis to those obtained when it is applied to the oscillators pairs within the Kuramoto model.

The MEG phase differences of the two motor cortices at rest showed long-range temporal correlations of their rate of change consistent with the pre fully synchronous state. In contrast finger movement was found to disrupt the phase difference fluctuations away from the LRTC regime. The principle of correlations breaking down has been observed previously in the amplitude fluctuations of MEG and EEG [15, 44]. The phase relationship of oscillatory neurophysiological signals is a measure of communication between neuronal pools [3, 12, 27, 45–47]. We suggest that LRTCs in the phase relationship of motor cortices show that their interaction at rest is in a state of optimal readiness. During movement, this interaction is disrupted because the cortices become engaged in activity. We further suggest that, in the active state, the cortices have a lower predisposition to synchrony.

Although our findings are related to previous studies of event related synchronisation and desynchronisation (ERS and ERD), e.g., [48, 49], our results are distinct in a number of ways. First, we remain agnostic about the synchronisation level within any specific neural region, but instead study the phase relationship between two brain regions, which is a property orthogonal to oscillation amplitude measures studied previously [24]. Second, our analysis is applied to time series that last several minutes, during continuous finger tapping rather than addressing the more transient effects of movement previously studied [6, 7].

It is important to note that DFA is dependent on finding power law scaling in the detrended fluctuation magnitude and that power law scaling in itself is not a sufficient condition for criticality [50]. We conclude that the interactions between motor brain regions at rest are indicative of a system in a regime that provides a suitable environment for the brain to react readily to external stimuli. Since we do not have access to the full spectrum of brain activity, we cannot actually say whether it is close to a critical transition. However, the ability of our methodology to identify a system with the potential to change could have important implications in the future for assessing the sensitivity of a human subject in responding to drugs or other treatments.

## Acknowledgements

Maria Botcharova was funded by the Centre for Mathematics and Physics in the Life Sciences and Experimental Biology (CoMPLEX), University College London. Simon F. Farmer was supported by University College London Hospitals Biomedical Research Centre (BRC), ’Moger Moves’ and the Szeben Peto fund. This work was supported by an MRC UK MEG Partnership Grant, MR/K005464/1. The WTCN is supported by the Wellcome Trust.

## A Tables

**Table 1:** P-values for one-tailed Student’s t-tests between the mean difference in DFA exponents between resting state and movement conditions against four sets of control data (white noise - Gauss, scanner noise - Scanner, phase-shuffled rest data - ShuffledR, phase-shuffled tapping data - ShuffledB). Significant tests are shown in bold. Only DFA exponents determined to be valid by ML-DFA are included in the calculation. The number of valid exponents for each set is provided in Table 2.

|  | Frequency Band (Hz) |  |  |  |  |  |
| --- | --- | --- | --- | --- | --- | --- |
| Data | 4-6 | 6-8 | 8-10 | 10-12 | 12-14 | 14-16 |
| Gauss | 0.14 | 0.24 | 0.29 | **0.02** | 0.13 | 0.25 |
| Scanner | 0.30 | 0.78 | 0.17 | 0.14 | **0.03** | 0.19 |
| ShuffledR | **0.01** | **0.01** | **0.01** | **0.01** | **0.01** | **0.02** |
| ShuffledB | **0.01** | **0.02** | **0.01** | **0.01** | **0.02** | **0.03** |
| Data | 15-17 | 17-19 | 19-21 | 21-23 | 23-25 | 25-27 |
| Gauss | 0.27 | 0.46 | 0.34 | 0.09 | **0.00** | 0.10 |
| Scanner | 0.19 | 0.22 | 0.05 | **0.04** | **0.00** | **0.03** |
| ShuffledR | **0.01** | **0.02** | **0.02** | **0.01** | **0.01** | **0.01** |
| ShuffledB | **0.02** | 0.07 | 0.10 | 0.11 | 0.26 | 0.05 |
| Data | 27-29 | 29-31 | 31-33 | 33-35 | 35-37 | 37-39 |
| Gauss | 0.28 | 0.11 | 0.07 | 0.66 | 0.15 | 0.15 |
| Scanner | 0.86 | 0.27 | 0.09 | 0.76 | 0.30 | 0.39 |
| ShuffledR | 0.11 | 0.05 | 0.08 | 0.27 | 0.14 | 0.18 |
| ShuffledB | 0.09 | 0.07 | 0.11 | 0.33 | 0.59 | 0.15 |

**Table 2:** Number of DFA exponents determined to be valid by ML-DFA for each of type of time series (labeling as in Appendix Table 1). Only valid exponents were used in calculating average DFA exponents.

|  | Frequency Band (Hz) |  |  |  |  |  |
| --- | --- | --- | --- | --- | --- | --- |
| Data | 4-6 | 6-8 | 8-10 | 10-12 | 12-14 | 14-16 |
| Rest | 2 | 2 | 6 | 5 | 6 | 6 |
| Both | 1 | 5 | 3 | 5 | 7 | 6 |
| Gauss | 3 | 2 | 6 | 4 | 3 | 6 |
| Scanner | 1 | 1 | 3 | 4 | 5 | 4 |
| ShuffledR | 97 | 95 | 98 | 97 | 95 | 96 |
| ShuffledB | 99 | 100 | 94 | 95 | 95 | 98 |
| Data | 15-17 | 17-19 | 19-21 | 21-23 | 23-25 | 25-27 |
| Rest | 5 | 5 | 5 | 7 | 6 | 7 |
| Both | 6 | 6 | 7 | 5 | 7 | 7 |
| Gauss | 6 | 4 | 7 | 5 | 7 | 4 |
| Scanner | 6 | 6 | 7 | 4 | 7 | 7 |
| ShuffledR | 97 | 96 | 97 | 94 | 94 | 95 |
| ShuffledB | 98 | 97 | 96 | 98 | 99 | 97 |
| Data | 27-29 | 29-31 | 31-33 | 33-35 | 35-37 | 37-39 |
| Rest | 6 | 7 | 5 | 7 | 6 | 6 |
| Both | 7 | 6 | 7 | 7 | 6 | 7 |
| Gauss | 6 | 7 | 7 | 7 | 7 | 6 |
| Scanner | 6 | 6 | 5 | 7 | 7 | 6 |
| ShuffledR | 97 | 93 | 96 | 95 | 96 | 95 |
| ShuffledB | 95 | 97 | 97 | 95 | 96 | 94 |

